# Cryopreservation of brain organoids - a tool for on-demand organoid banking

**DOI:** 10.64898/2026.05.19.726365

**Authors:** Lixuan Ding, Jingyi Zhang, Dowlette-Mary Alam El Din, Itzy E. Morales Pantoja, Thomas Hartung, Lena Smirnova

## Abstract

Cryopreservation offers an option for long-term storage and global distribution of complex *in vitro* models, yet protocols for multicellular microphysiolgocial systems (MPS) such as brain organoids/spheroids remain limited. Here, we systematically compared three commercially available cryopreservation (mFreSR, CryoStorCS10, and 3dGRO) and two freezing time points, and established a robust workflow for freezing and recovering brain organoids. After defrosting, we assessed morphology and metabolic activity. We also evaluated electrophysiology, calcium transients, and neurite outgrowth. In addition, we measured astrocyte migration, apoptosis, mitochondrial integrity, microglia survival, and neural marker expression. We found that organoids require a 4-week recovery period to regain structural and functional stability. Although organoids frozen at week 6 showed higher metabolic activity after recovery, organoids cryopreserved at week 2 had clearly better functional outcomes. They exhibited stronger spontaneous network firing and maintained calcium transients. Finally, incorporated microglia-like cells survived the freezing and displayed comparable morphology to unfrozen controls. Across the endpoints measured here, 3dGRO showed the most favorable overall performance; formal ranking across media awaits harmonized normalization, single-organoid electrophysiology, and prespecified QC thresholds. Together, these results define a practical and reproducible cryopreservation strategy that preserves key physiological features of brain organoids and supports the establishment of ready-to-use organoid banks. The ability to reliably store and distribute complex brain-like tissues represents an essential step toward global standardization, scalable experimentation, and wider adoption of human-relevant microphysiological systems. Together, these results demonstrate recovery of key physiological features in the subset of organoids that remain viable after thaw and support the feasibility of brain organoid banking.

## INTRODUCTION

Cryopreservation is a critical technique in biological research and clinical applications, enabling the long-term storage of biological materials by halting metabolic and biochemical processes and effectively preserving their structural and functional integrity ^1^. Cryopreservation of cells has been well-established, facilitating their use in various experimental and therapeutic contexts and enabling large-scale banking, storage, and distribution worldwide through cell banks or individual researchers, fostering broad collaborations and harmonization of research ^2^. Cryopreservation of tissues and recently emerging Microphysiological Systems (MPS, such as organoids and spheroids)^3–6^, however, is less developed, especially for preserving very complex multicellular structures. The challenges include preventing ice crystal formation, which can cause structural and mechanical damage to tissue and cellular integrity, and maintaining cell viability and functionality upon defrosting^7,8^. In a larger 3D construct, temperature gradients will unavoidably differ between core and periphery, challenging the goal of defined freezing protocols. The complexity of brain organoids, which comprise diverse neuronal populations and glial cells, adds another layer of difficulty; different cell types vary in their sensitivity to cryopreservation protocols, and the intricate neuron-neuron and neuron-glia networks and circuitry must be re-established after thawing.

Recent advancements in the field have demonstrated the utility of brain organoids for studying various neurological diseases and for drug and chemical testing^9–12^. Cryopreservation of brain organoids derived from different iPSC lines with diverse genetic backgrounds, derived from various individuals and patients, offers significant opportunities for the biomedical field. This technique allows banking of organoids at different stages of differentiation, ensuring standardized differentiation protocols and facilitating the dissemination of pre-differentiated organoids globally.

Cryopreservation involves the preservation of tissues, cells, organelles, proteins, scaffold by cooling them to extremely low temperatures ^1^. However, the process is sensitive, often resulting in 20-30% cell death due to osmotic stress, membrane damage, and ice crystal formation ^13^. Despite these challenges, several recent studies have successfully cryopreserved various types of organoids, including hepatic, intestinal, and colon organoids, and, very recently, the first proof of the possibility of cryopreserving cortical organoids was published^14–17^.

Studies over the last three years have shown that brain organoids derived from iPSCs, hPSCs, and primary tissue can be successfully cryopreserved using several approaches, including conventional slow cooling with DMSO-based solutions, vitrification adapted from embryo preservation techniques, a novel MEDY cocktail incorporating methylcellulose, ethylene glycol, DMSO, and the ROCK inhibitor Y27632, and slow freezing on an integrated pillar plate platform. Several studies were identified addressing the cryopreservation of human brain organoids, spanning from 2023 to 2025. The studies encompass a range of organoid types, cryopreservation techniques, and source materials. One resource optimized a slow-freeze protocol for iPSC-derived brain organoids and chondrospheres^18^. Two sources^16,17^ describe the same MEDY method, with the latter providing a detailed protocol. Similarly, two sources from^19,20^ describe the same pillar plate cryopreservation platform, which is distinctive in that it integrates cryopreservation, thawing, culturing, staining, and imaging within a single system. Several consistent findings emerge. First, all studies employing iPSC-derived organoids with modern protocols reported preservation of functional markers post-thaw, though with varying degrees of quantitative fidelity. Zolfaghar et al. noted that while early progenitor markers (OCT4, PAX6, FOXG1) were fully preserved, mature neuronal markers (MAP2, CTIP2, TUBB3) were slightly reduced relative to controls, which is consistent with cryopreservation at early developmental stages where these mature markers are less established. Ramani et al. reported notably high recovery rates of 75–90% for 18-day-old Hi-Q organoids, whereas organoids frozen at later stages (days 35 or 40) showed damaged cytoarchitecture and elevated apoptosis and could not be successfully recultured^21^. Both Mojica-Perez et al. and Ramani et al. confirmed that electrical activity and mature neuronal markers were preserved post-thaw, providing critical evidence that cryopreserved organoids retain functional relevance for electrophysiological and neurodevelopmental studies^21,22^.

Immune-competent, microglia-containing brain organoids could be another challenge for cryopreservation. Microglia are highly responsive immune cells which can rapidly change their phenotype in response to environmental stress, including osmotic imbalance, thawing-associated injury, and cellular debris release from damaged cells^23,24^. Unlike the assessment of neurons and astrocytes, which often focuses on survival, morphology, and functional recovery, the evaluation of microglia-containing brain organoids also determines whether microglia preserve an appropriate homeostasis, since microglia are highly plastic immune cells that can rapidly shift their phenotype in response to cellular stress and debris during the freezing and recovery process^25^. We previously successfully preserved and banked microglia progenitors^26^; however, the recovery upon defrosting was better at the stage of hematopoietic cells than microglia progenitors or mature microglia. Therefore, we examined the microglia survival within the frozen organoids as the most vulnerable population. Hence, establishing reliable cryopreservation strategies for these more complex organoids would further expand the utility of brain organoids by enabling standardized and on-demand models that preserve not only neural architecture and electrophysiological, but also immune component of the human brain microenvironment.

The apparent variation in cryopreservation success across studies can be largely reconciled by considering three key variables: organoid size/stage, CPA formulation, and the cooling method employed. There is no universal method yet and each and every model has to be optimized. In the case of our model, the availability of cryopreservation has several impacts: creation of an organoid bank with different genetic backgrounds, which can then be distributed across the co-elaborators. This model is also commercially available via 28.bio and can be provided on demand to end users.

In this study, we established a method for the cryopreservation of our brain MPS by using a commercially available product designed for long-term preservation. We extensively characterized the cultures’ viability and functionality upon defrosting. Although our protocol allows successful preservation of the main cell types in brain organoids, cryopreservation at earlier stages of differentiation and a recovery period for up to 4 weeks are advisable to ensure high viability, functionality and the physiological ratio of neurons and glial cells. In agreement with previous studies, we confirmed that it is advisable to cryopreserve at earlier stages, with a better recovery rate than at more mature stages, but both stages can be re-cultured.

The ability to cryopreserve brain organoids including immune-competent organoids is an advancement in the field that enables the creation of banks of disease models accessible on demand, significantly enhancing interlaboratory reproducibility and accelerating research in neurobiology and pharmacology.

## RESULTS

### Brain organoids recover after defrosting

To pursue our goal of developing reliable freezing conditions, we selected three freezing media: mFreSR (StemCell Technologies), CrystorCS10 (Biolife Solutions®), and 3dGRO (Sigma-Aldrich). To investigate whether the differentiation stage at the time of freezing affects survival and recovery, organoids were cryopreserved at two time points: week 2 (W2, 2-week group) and week 6 (W6, 6-week group) of differentiation. Post-thaw recovery was assessed after 24 hours and after 2 and 4 weeks in culture (Figure 1). Organoids, which remained for the entire time in the incubator, served as a control. A substantial amount of cellular debris and irregularly shaped organoids were observed 24 hours after thawing. In contrast, organoids exhibited better structural integrity (round shape and no debris) after 4 weeks of recovery at both freezing time points (Figure S1A-B, and S2A). Because this assessment was based on bright-field morphology and 2D diameter, internal cytoarchitecture was not quantitatively assessed. In subsequent experiments, we assessed organoid size, viability, neurite outgrowth, glial migration, electrical activity, mitochondrial membrane potential, cleaved Caspase3, and expression of key neural markers.

**Figure 1.**
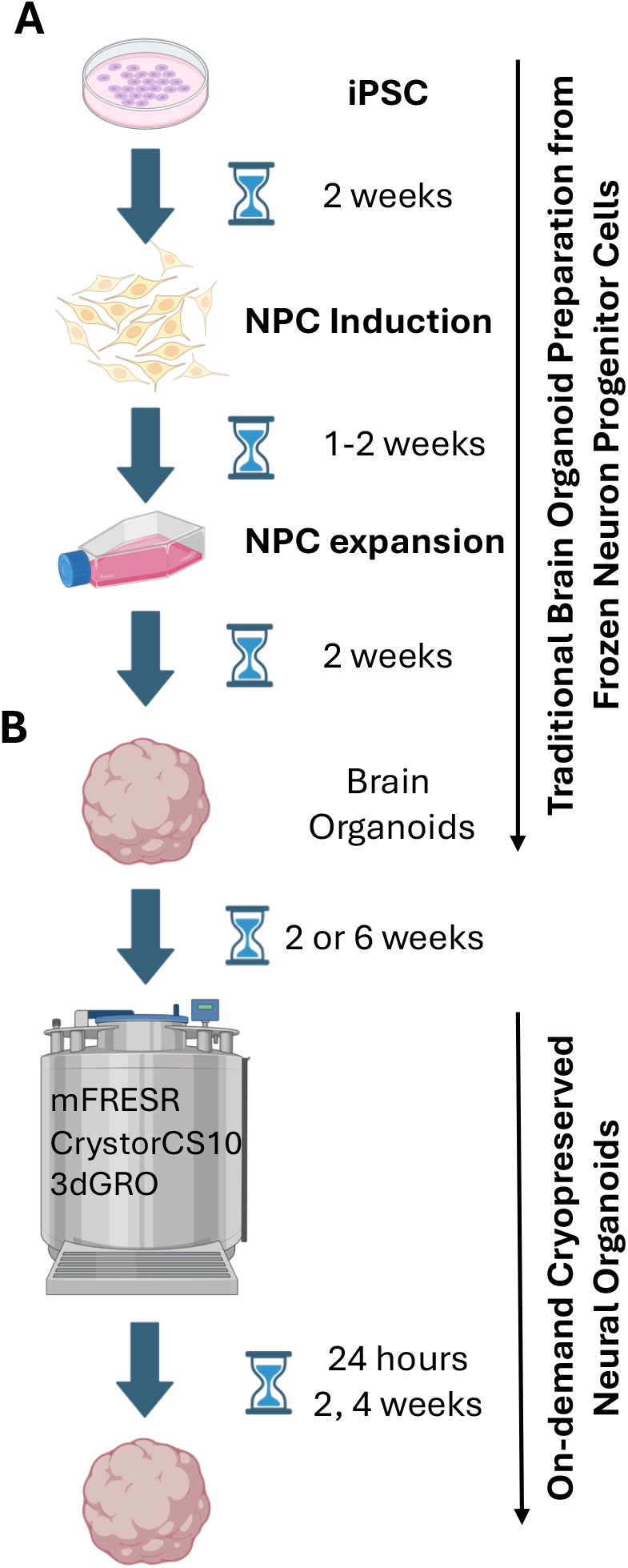
Experimental workflow of brain organoid generation and cryopreservation. Schematic of the process and time before testing for (A) preparing a brain organoid from frozen stocks of c and (B) taking a brain organoid directly from frozen stocks.

### Frozen at week 2 or 6, organoids were smaller than controls, with better recovery when frozen at week 2

To quantify average organoid size after defrosting, we randomly selected three fields per condition and measured at least 15 organoids per field. At least 40 organoids were assessed per condition/time point. The size of freshly thawed organoids (24 hours post-defrosting) was very similar to the unfrozen control condition, but decreased during the first 2 weeks of recovery. Although the size increased at 4-weeks, overall, frozen organoids remained smaller than control organoids even after a 4-week recovery period (Figure 2, S2B). Organoids frozen in CryostorCS10 medium at 6 weeks remained the smallest, while organoids frozen in mFreSR medium were the closest to the control size at 4 weeks post-recovery. These changes in size across conditions emphasize the need for a recovery period to stabilize the cultures.

**Figure 2.**
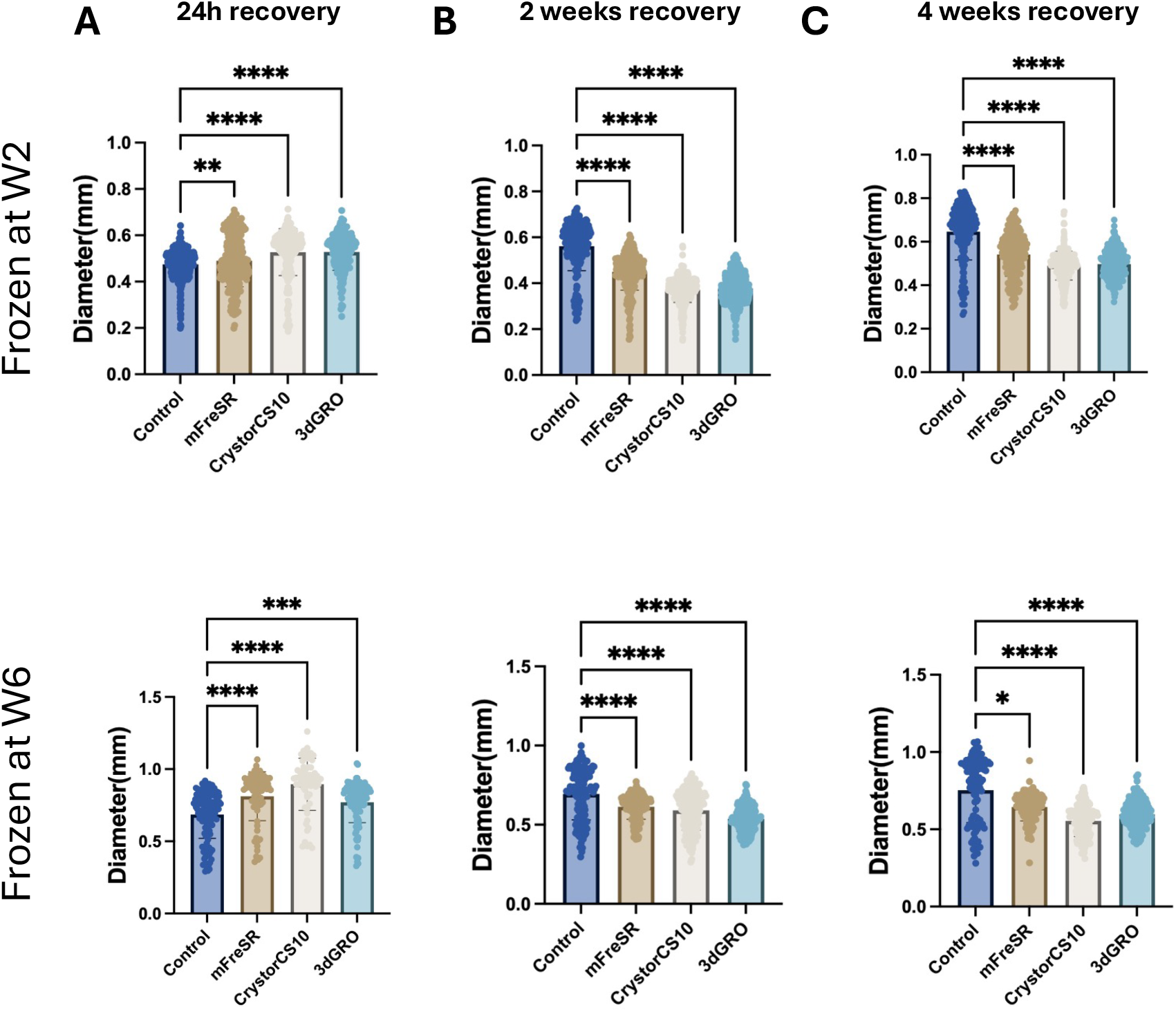
Size quantification of cryopreserved brain organoids. Organoids were frozen at 2 weeks (W2) or 6 weeks (W6) and recovered for 24 hours, 2 or 4 weeks. Organoid diameter was measured at (A) 24h, (B) 2 weeks, and (C) 4 weeks after thawing. Age-matched organoids maintained continuously in the incubator were used as controls. Data are presented as Mean ±SD from 3 biological replicates with at least 40 organoids measured in total for each condition and time point. One-way ANOVA with Dunnett’s post-hoc test was used to assess statistical significance. * p<0.05, ** p<0.01, ****p<0.0001

### Metabolic activity was different immediately after defrosting and after recovery

The Resazurin Reduction assay was performed to assess the viability and metabolic activity of cryopreserved brain organoids compared with age-matched unfrozen controls. To account for the size difference, viability/metabolic activity data were normalized to an average organoid size within each group. Metabolic activity was significantly reduced 24 hours after defrosting (Figure 3A) and was higher than in controls two weeks after recovery (Figure 3B). Four weeks post-thaw, organoids frozen at week 6 of differentiation showed significantly higher metabolic activity than controls, whereas the increase of metabolic activity in organoids frozen at 2 weeks was significant but to a lesser extent (Figure 3C). Because resazurin reflects aggregate reducing capacity rather than per-cell metabolic flux, these data do not distinguish increased per-cell metabolism from altered cell number or cellular composition. Size and viability data suggest that the younger organoid group (frozen at 2 weeks) recovers better than older organoids (frozen at 6 weeks), and a 4-week recovery period is ideal. Thus, we focused on this time point in subsequent experiments.

**Figure 3.**
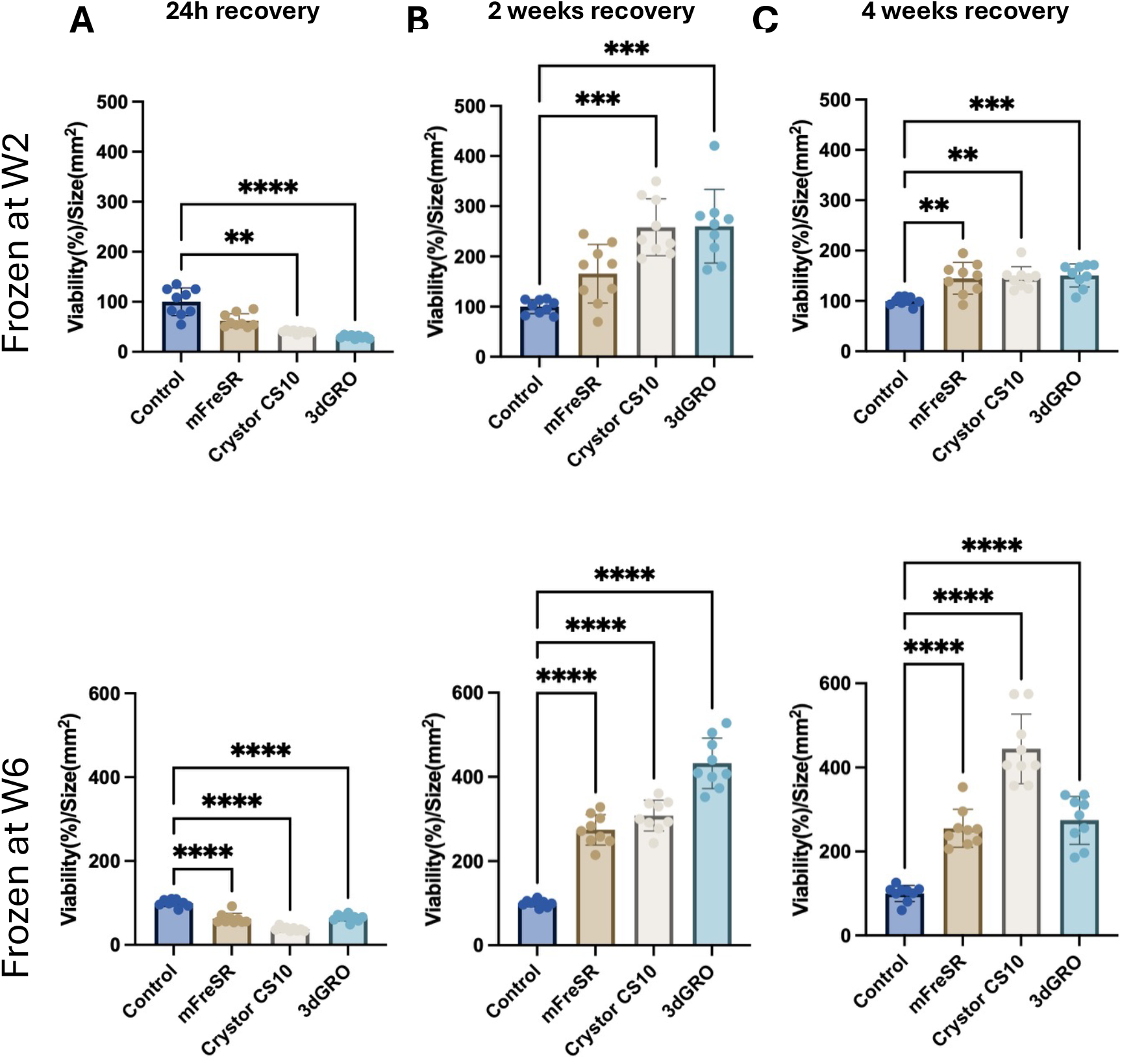
Percentage of metabolically active cells in cryopreserved brain organoids. Organoids were frozen at 2 or 6 weeks. Viability was normalized to the average organoid area and corresponding age-matched incubator controls. Viability was assessed with the Resazurin Reduction Assay at (A) 24h, (B) 2 weeks, and (C) 4 weeks after recovery. Data are presented as Mean ±SD from 3 biological replicates with three technical replicates (10 organoids per replicate). Kruskal-Wallis test with Dunn’s post-hoc test was used to assess statistical significance. ** p< 0.01, *** p< 0.001, **** p< 0.00001.

### Spontaneous electrical activity was preserved after defrosting

To further evaluate the functionality of the frozen cultures, we assessed spontaneous electrical activity using a high-density microelectrode array (HD-MEA) system and calcium imaging.

For HD-MEA, frozen at week 2 (Figure 4A-B, 2-week group) and week 6 (Figure 4C-D, 6-week group) organoids were in recovery period for 4 weeks, then plated on HD-MEA, and spontaneous electrical activity was recorded over time. We analyzed the parameters related to spontaneous bursting and spiking activity in brain organoids, including burst frequency, spikes within bursts, mean spikes per burst, mean spikes per burst per electrode, mean burst duration, mean burst peak firing rate, mean interburst interval, mean interspike interval within bursts, interspike interval outside bursts, and spike mean firing rate. Representative active area and raster plots show both frozen and control organoids (Figure 4A (frozen at week 2) and 4C (frozen at week 6)) demonstrate spontaneous spiking and bursting dynamics. In addition, spontaneous bursting and spiking activity were quantified over time across different groups to determine whether spontaneous electrical activity was maintained in frozen organoids (Figure 4B and 4D).

**Figure 4.**
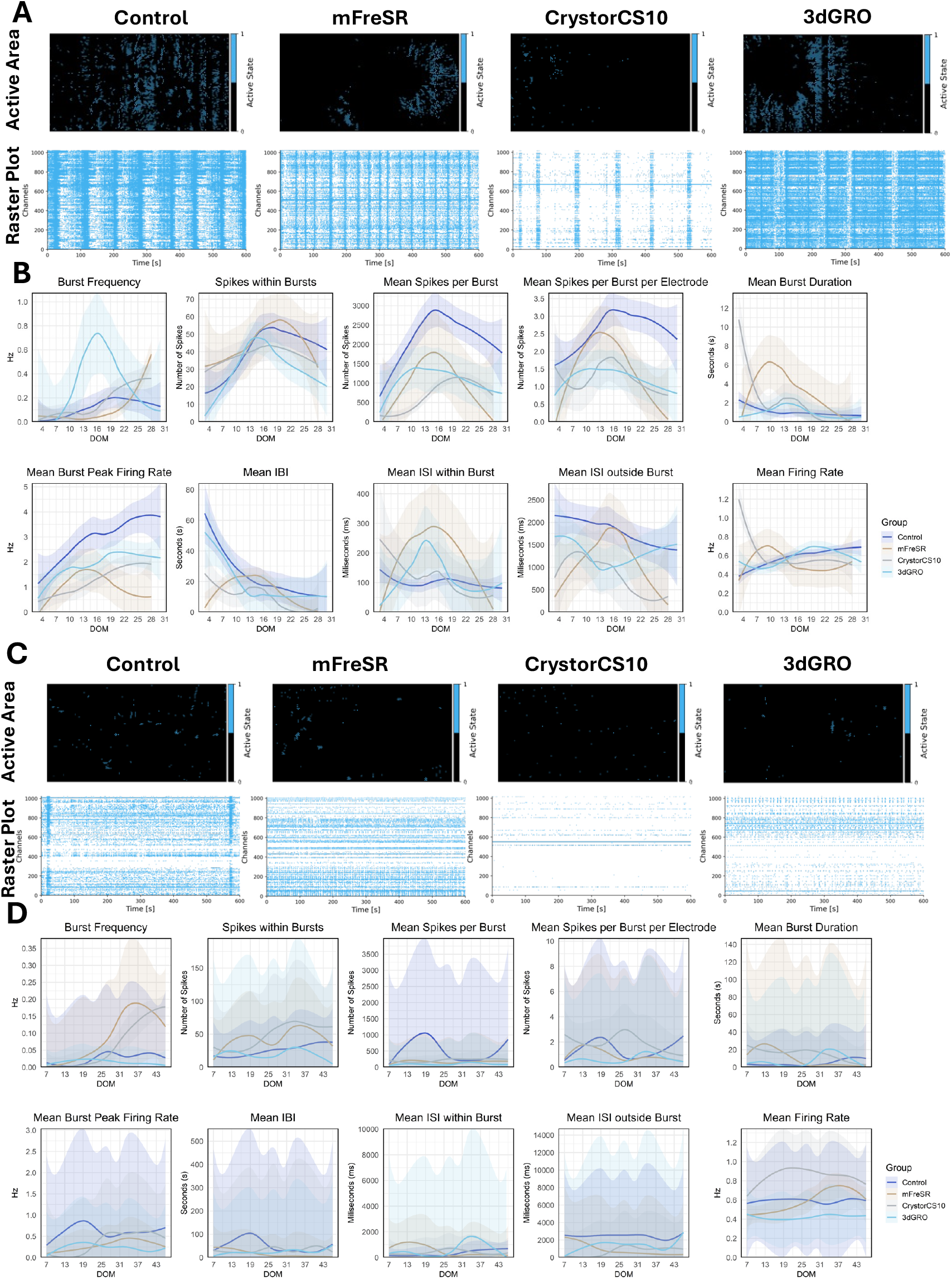
Assessment of electrical activity in organoids after cryopreservation at 2 and 6 weeks and 4 weeks of recovery. A) Representative raster plots show spontaneous synchronized firing events in cryopreserved and incubator control organoids. B) Network activity metrics from all organoid groups over time (blue line represents incubator control organoids, brown line represents mFreSR, grey line represents CrystorCS10, and the light blue line represents 3dGRO medium, respectively). The darker lines shown represent the mean, and the shaded region represents the standard deviation. The data shown are plotted from at least three wells per group and time point across two independent experiments. ISI: Interspike Interval. IBI: Interburst Interval. DOM: Days on MEAs.**C**) Representative raster plots show spontaneous synchronized firing events in cryopreserved incubator control organoids. D) Network activity metrics from all organoid groups over time (blue line represents control organoids, brown line represents mFreSR, grey line represents CrystorCS10, and the light blue line represents 3dGRO medium, respectively). The darker lines shown represent the mean, and the shaded region represents the standard deviation. Data shown are plotted from at least three wells per group and timepoint across one independent experiment. ISI (Interspike Interval). IBI: Interburst Interval. DOM: Days on MEAs.

For both 2- and 6-week groups, all conditions showed organoid attachment to the HD-MEA surface and exhibited spontaneous spiking and bursting as shown in the representative raster plot (Figure 4A and 4C). When quantified, all groups showed quantifiable spontaneous activity and showed similar trends over time (Figure 4B and 4D). Overall activity, however, was lower at the 6-week time point for all samples, including unfrozen control, when compared to the young group. Notably, there was a high variability among organoids in the 6-week group across all samples, as shown by the shaded error area (Figure 4D). Higher standard deviation and lower bursting and spiking frequency were consistent with our previous observations of organoids at this age (10 weeks in culture).

Together, these results indicate that brain organoids retain functional spontaneous electrical activity from neuronal networks after cryopreservation at both 2 and 6 weeks and a 4-week recovery period, with overall lower activity in the older group.

To confirm our findings from the HD-MEA analysis, we used calcium imaging to quantify spontaneous electrical activity. Thus, we employed Fluo-4, a fluorescent calcium indicator dye, to monitor spontaneous calcium fluxes and calculate the change in fluorescence intensity (ΔF/F). For the 2-week group, rise time, decay time, peak amplitude, and burst duration did not differ significantly between control organoids and those preserved in mFreSR or 3dGRO. Although mFreSR increased the number of detectable peaks, it did not affect peak waveform characteristics. In contrast, no active organoids were detected in the CryoSTOR-frozen group (Figure 5A). We then left organoids to recover for additional 2 weeks and assessed the Ca^2+^ transients after a total of 6 weeks of recovery and found that all organoids in all groups were active (Figure 5B). 3dGRO significantly increased decay time, peak amplitude, and burst duration, while decreasing the number of peaks. No significant differences were observed in the mFreSR or CrySTOR-frozen groups. Lastly, no spontaneous calcium transients were detected in week-6 organoids under the present imaging conditions and regardless of the medium used (Figure 5C). Results showed that calcium signaling activity was preserved in the 2-week group after 4 and 6 weeks post-defrosting, but not in the 6-week group. In week-6 organoids, HD-MEA spikes were detected whereas spontaneous calcium transients were not detected, consistent with modality-specific sensitivity in dense 3D tissue. Pharmacological validation of MEA spikes and evoked calcium controls were not performed in this study.

**Figure 5.**
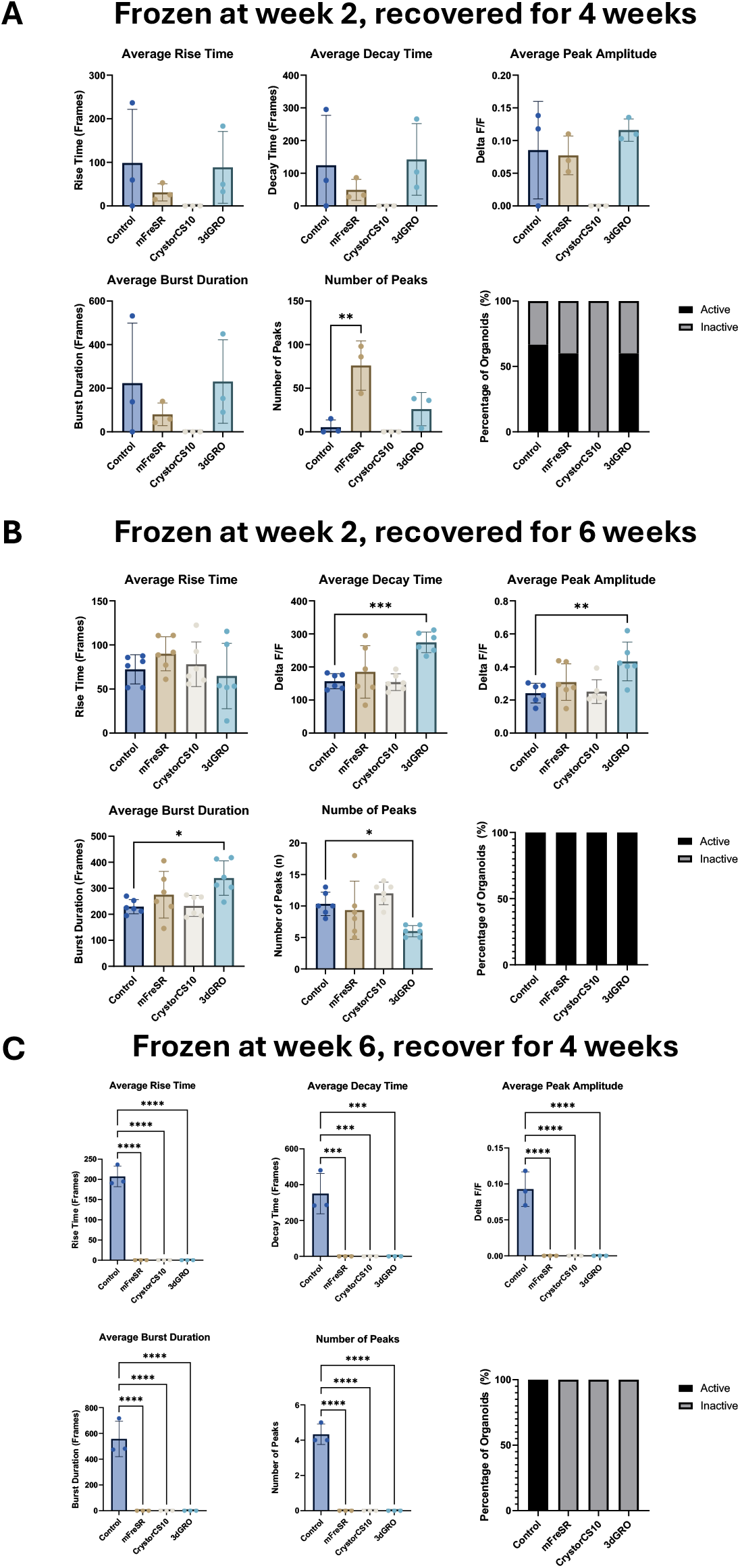
Assessment of Ca^2+^ flux in organoids after cryopreservation and recovery. Measurements of rise time, decay time, peak amplitude, burst duration, and the number of calcium peaks were performed on brain organoids cryopreserved at 2 or 6 weeks of differentiation and recovered for 4/6 weeks. Results showed that the Calcium signaling activity was preserved in the 2-week group but not in the 6-week group. Statistics were calculated using a one-way ANOVA with Dunnett’s multiple comparisons test. p < 0.05 was considered significant. For organoids frozen at week 2 and recovered for 4 weeks, calcium imaging was performed on one independent experiment with at least 3 organoids per group. For organoids frozen at week 2 and recovered for 6 weeks, calcium imaging was performed on one independent experiment with at least 6 organoids per group. For organoids frozen at week 6 and recovered for 4 weeks, calcium imaging was performed on one independent experiment with at least 3 organoids per group.

Since the organoids frozen at week 2 showed functionality more comparable to that of age-matched controls, we prioritized the week 2 cryopreservation group for the subsequent experiment.

### The neurite outgrowth was enhanced after defrosting, whereas extension of GFAP-positive processes was unchanged

Neurite elongation and outgrowth were used as functional endpoints for assessing the impact of cryopreservation on neurons and neurogenesis. The GFAP Sholl endpoint used here reflects astrocytic process extension relative to the organoid edge rather than direct soma migration. GFAP-positive process extension / Sholl analysis was evaluated in parallel to analyze astrocyte functionality. No statistically significant differences were observed in neurite outgrowth and astrocyte migration at the 6-week time point (Supplementary Figure S3). In contrast, neurite outgrowth was significantly increased in organoids frozen at week 2, which correlates with their smaller size and higher metabolic activity (Figure 6A). Interestingly, organoids frozen in 3dGRO medium showed an incremental increase and most closely matched the neurite outgrowth in the incubator controls. In agreement with this observation, we previously reported faster and more extensive neurite outgrowth in organoids with a smaller diameter^27^. No statistically significant changes in glia migration were observed across all groups at the 2-week freezing time point (Figure 6B).

**Figure 6.**
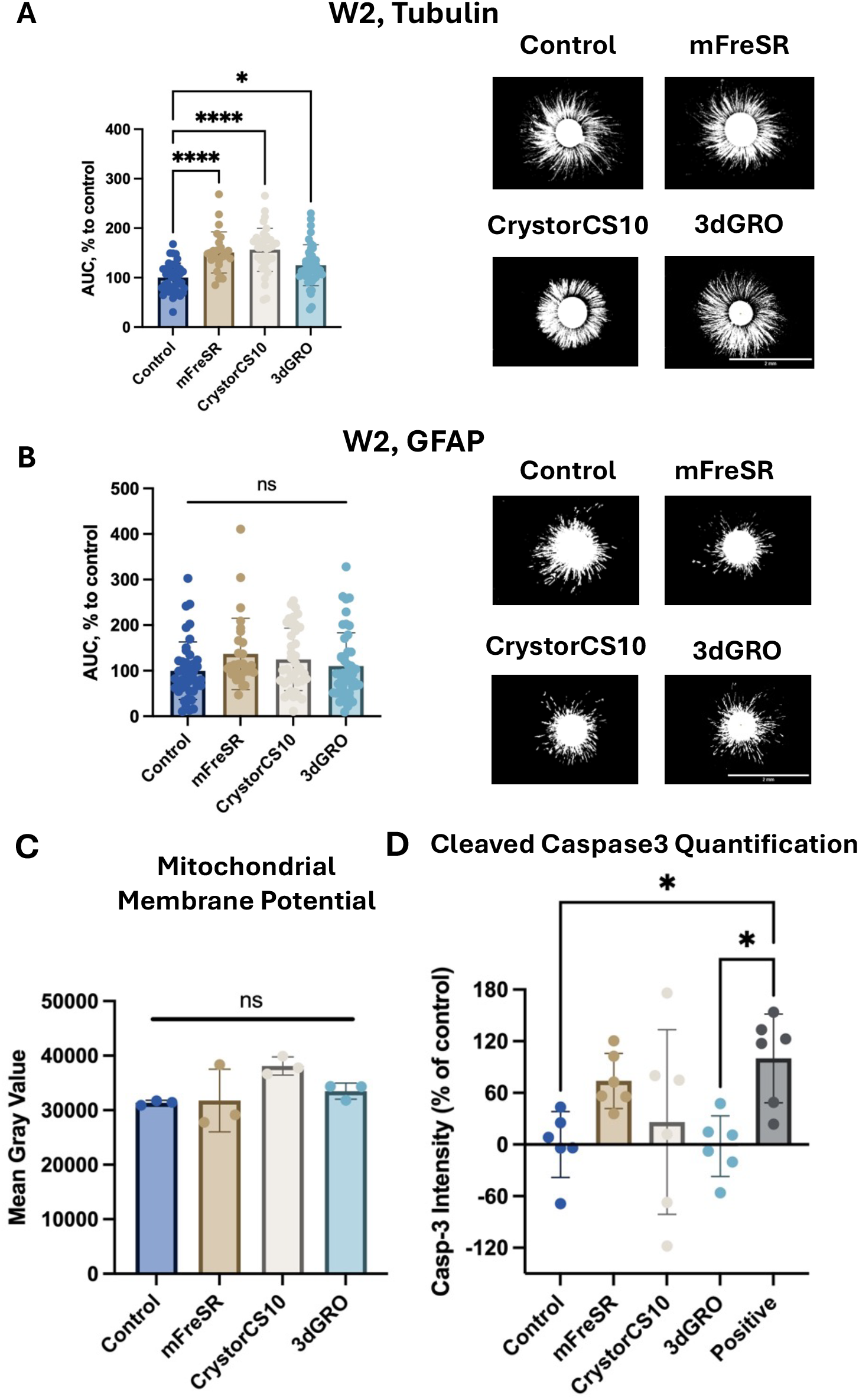
Quantification of neurite outgrowth, astrocyte migration, mitochondrial membrane potential, and cleaved Caspase 3 in brain organoids after cryopreservation at 2 weeks and 4 weeks of recovery. Organoids were frozen at week 2 (W2) and recovered for 4 weeks. Area Under the Curve (AUC) from Sholl analysis of (A) neurite outgrowth (β-III-Tubulin) and (B) astrocyte migration (GFAP) is shown. The area under the curve (AUC) was calculated for each condition, and each graph represents the Mean ± SD from 3-18 organoids per independent experiment, with five independent experiments for β-III-Tubulin and three - for GFAP. Kruskal-Wallis test with Dunn’s post-hoc test was used to assess statistical significance. (C) Mitochondrial membrane potential, quantified by mean gray value. Data represent Mean ± SD from 3 organoids per condition. Statistical significance was assessed with Kruskal-Wallis test and Dunn’s post-hoc test. ns – not significant. (D) Quantitative analysis of overall intensity of cleaved Caspase 3 staining, normalized to negative and positive (Staurosporine) controls. Data represent mean ± SD; n = 6 organoids per condition from two independent experiments. Significance was assessed with Kruskal-Wallis test with Dunn’s post-hoc test. *p<0.05, ***p<0.001

### No changes in mitochondrial membrane potential and no increased apoptosis in 2-week frozen organoids after 4 weeks of recovery

To further evaluate organoid recovery after cryopreservation at 2 weeks and 4-week recovery, mitochondrial membrane potential was quantified with the MitoTracker Assay, resulting in no significant difference across the groups (Figure 6C, and Suppl Figure S4).

We assessed levels of cleaved Caspase-3 as an apoptosis marker in the group cryopreserved at week 2 and recovered for 4 weeks ^28^. Quantitative analysis of immunofluorescence staining revealed no significant difference in cleaved caspase-3 intensity between cryopreserved organoids and age-matched unfrozen incubator controls after 4 weeks of recovery (Figure 6C and Suppl. Figure S5). A positive control (Staurosporine-exposed organoids), showing a higher level of apoptosis, confirmed the specificity and sensitivity of the assay. Of note, the level of cleaved Caspase 3 in organoids frozen in 3dGRO was most comparable with the incubator control, and these were the two conditions that were significantly different from the positive control. Taken together, the Caspase-3, neurite outgrowth, and electrophysiology results suggest that 3dGRO provides improved cryoprotection. These organoid-level averages, however, do not resolve spatial or cell-type-specific heterogeneity.

### The cellular composition was preserved in 2-week frozen organoids after 4 weeks of recovery

To evaluate whether the cryopreservation affected the cellular composition of brain organoids, we performed immunofluorescence staining and gene expression analysis in 2-week group organoids, which were recovered for 4 weeks after defrosting. Confocal imaging showed preserved structural organization of neurons and synapses across all conditions, with β-tubulin (neuronal cytoskeleton), MAP2 (neuronal dendrites), and synapsin (presynaptic marker) staining comparable between frozen and control organoids (Figure 7A).

**Figure 7.**
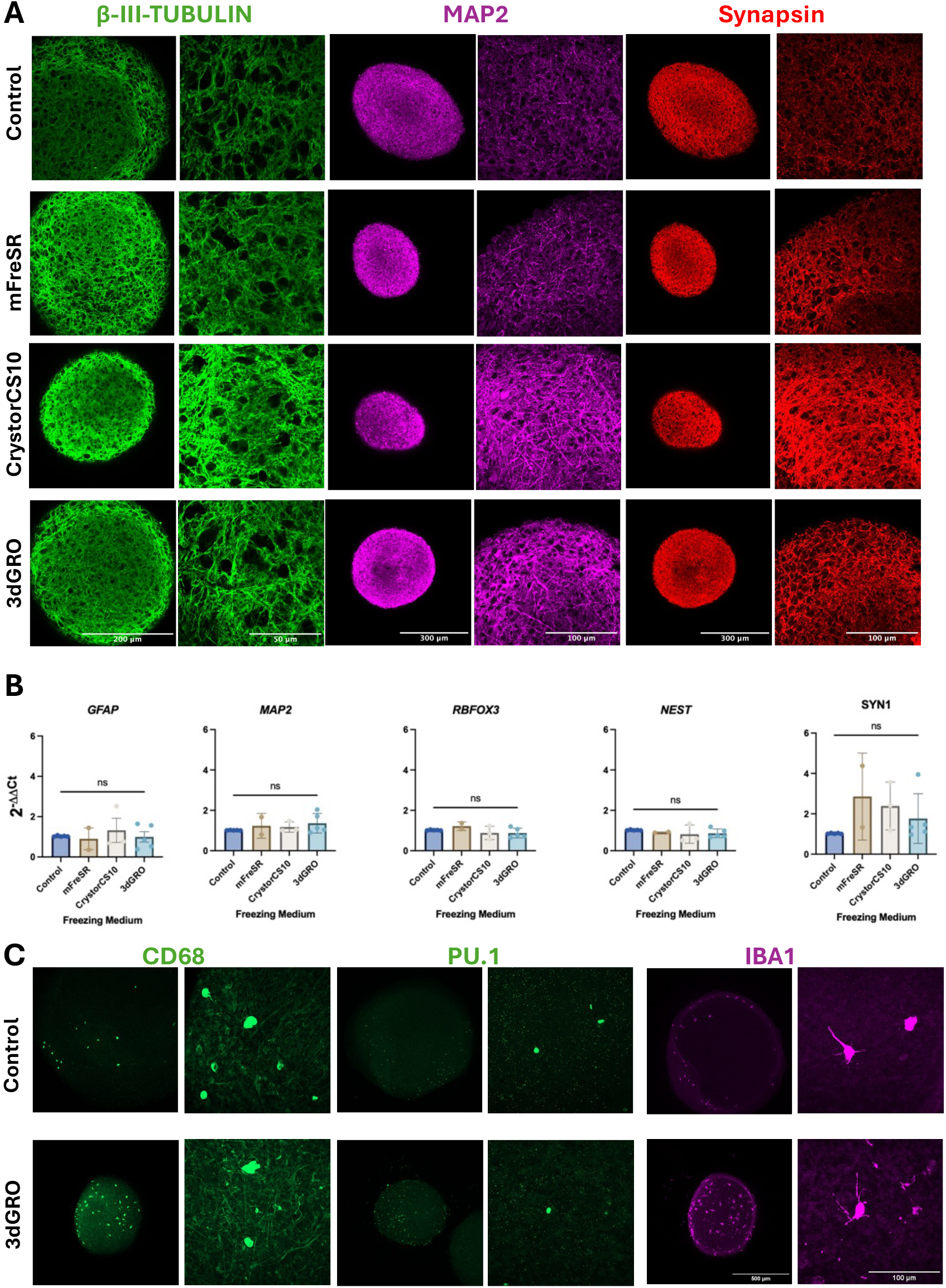
Expression of main brain markers in brain organoids after *3Dgro* cryopreservation and recovery. Organoids were frozen at week 2 (W2) of differentiation and recovered for 4 weeks. Age-matched organoids that remained in the incubator throughout the experiment served as a control. (A). Representative confocal microscopy images of brain organoids stained for β-tubulin (green), MAP2 (magenta), and Synapsin (red). (B) Quantitative RT-PCR analysis shows no significant differences in the expression of *MAP2, NEUN, GFAP, NESTIN*, and *SYN1* between frozen and control organoids after 4 weeks of recovery. Data represent mean ± SD from two to five independent experiments with three biological replicates per experiment. For 3dGRO and control, five independent experiments were analyzed, while for CrystorCS10, two independent experiments were used. Statistical significance was calculated with One-way ANOVA. ns – not significant. (C) Preservation of microglia in brain organoids. Representative confocal microscopy images of age-matched control and 3dGRO-frozen organoids stained for microglia-associated markers CD68 (green), PU.1 (green), and IBA1 (purple).

The observation was further confirmed by RT-PCR in frozen organoids vs. age-matched unfrozen control organoids. We tested the expression of *MAP2* (marker for neuronal dendrites), *NEUN* (or *RBFOX3* marker for postmitotic neurons), *GFAP* (astrocytic marker), *NEST* (marker of neural progenitor cells), and *SYN1* (synaptic marker). Our results showed no significant difference in the expression of these markers between the two groups (Figure 7B). Although *SYN1* was slightly higher in the frozen organoids, consistent with the immunostaining, this difference did not reach statistical significance. These data support preservation of the measured neuronal and astrocytic markers after freezing at week 2 of differentiation and 4-week recovery. Overall, these results indicate that the major cellular populations were preserved following cryopreservation.

In addition to the neuronal and astrocytic markers, we also assessed whether microglial populations were preserved after cryopreservation of the immune-competent organoids. Immune-cell progenitors were incorporated into the organoids, and we followed the same protocol of cryopreservation and recovery. Confocal imaging showed that microglia-associated markers including CD68, PU.1, and IBA1 were detectable in frozen organoids after 4-week recovery and showed comparable distribution and morphology to age-matched unfrozen controls (Figure 7C). These microglial markers were observed together with preserved neural markers, including β-tubulin, MAP2, synapsin and astrocytic marker, GFAP (Supplemental Figure S6), suggesting that cryopreservation didn’t visibly disrupt the coexistence of microglial and neuronal populations within the organoid structure. Together, these results support the preservation of major neural, astrocytic, and microglial cellular components following cryopreservation.

## DISCUSSION

Previously, we developed a human-induced pluripotent stem cell (iPSC)-derived brain Microphysiological system with physiological levels of neuronal and glial cell populations ^29–31^. We recently incorporated microglia into these organoids, generating immune-competent organoids with long-term shelf-life^26^. The protocol allows production of large quantities of organoids: up to 4000 in one 6-well plate. Mass production allows scaling up and generating organoids on demand with diverse genetic backgrounds, including those derived from patients with the disease of interest. In addition, the reproducibility of iPSC-derived models remains a broadly recognized challenge and requires standardization. Thus, providing pre-differentiated organoids on demand will facilitate broader use of MPS models and drive international collaborations and harmonization efforts.

The challenge of organoid cryopreservation lies in their cell density, high-complexity structure, and heterogeneity of cellular composition ^32,33^. Beyond prior studies showing that mostly early-stage brain organoids can survive cryopreservation, this work establishes a standardized, commercially accessible workflow for a glia-containing neural MPS and defines, through multimodal post-thaw functional phenotyping, which freezing stage, medium, and recovery interval preserve neuronal network and neuron–glia-associated phenotypes most robustly. The novelty is that this is not mainly a new cryopreservation chemistry paper; it is a rigorous functional validation and workflow-standardization paper for bankable brain organoid MPS. The study tests three commercial media (mFreSR, CryoStorCS10, 3dGRO), two freezing stages (week 2 and week 6), and a defined recovery workflow in the same organoid system with the same assay framework. It includes broader post-thaw phenotyping and for the first time extends into a glia-containing brain MPS rather than mostly early neuroectodermal/cortical organoid settings. That is a real advance in comparability and practical standardization. It includes an explicit recovery-window recommendation, valuable boundary-setting result on older organoids and a pragmatic, adoption-ready workflow using accessible commercial media.

Therefore, our primary objective was to develop and validate a reproducible cryopreservation method for brain organoids. Key variables, such as the type of commercially available freezing media and the organoids’ recovery duration after defrosting, were modified based on previously tested cryopreservation techniques for other organoids. In this study, mFreSR, CrystorCS10, and 3dGRO freezing media have been evaluated in a set of endpoints to assess the integrity and functionality of the organoids upon defrosting.

The culture and differentiation of iPSCs, as well as organoid production, are labor-intensive processes that require the respective expertise and skills. The protocols are lengthy, spread over several months, and require precision. Developing a cryopreservation method enables the storage of numerous brain organoids from various donors, thereby reducing the time-consuming, ongoing production and maintenance. Although protocols vary, typically, it takes at least 4 weeks to generate and expand neural progenitors from iPSCs and another 8-20 weeks to get relatively mature organoids ready for use. The maturation step varies across protocols and researchers’ needs, with some organoids cultured for upto 6-9 months ^34–36^ or even years^37^. The differentiation protocol described here requires 8-9 weeks of maturation in 3D to obtain stable electrically active neural cultures with mature astrocytes and oligodendrocytes ^31^. Thus, cryopreservation can shorten the culture period by at least six weeks, eliminate the need for the laboratory to culture iPSC and NPCs routinely, and improve standardization. Even though a recovery period was needed, organoids retained the main cell types and functionality upon defrosting. In addition to neuronal and astrocytic populations, the ability to cryopreserve microglia within the organoids is particularly important because microglia represent the resident immune component of the central nervous system, play a key role in neural development, synaptic pruning, neuroinflammation, and, thus, is highly sensitive to environmental stress. This suggests that freezing at week 2 followed by a 4-week recovery period didn’t eliminate microglia-containing populations or grossly disrupt the association with neuronal structure. The organoid size quantification showed overall diameter decreases at 2 weeks after defrosting and was continuously recovering further during differentiation. Frozen organoids remained smaller than the control group, likely due to initial cell death during defrosting (as evidenced by debris in the first days of culture). The viability/metabolic activity was decreased significantly immediately after defrosting, suggesting cell death and lower metabolic activity in converting resazurin to resorufin. Interestingly, after 2 and 4 weeks of recovery, metabolic activity not only improved but was even significantly higher than the incubator control, when normalized to the size of the organoids, suggesting that remaining cells compensated for the stress with higher metabolic activity and, based on the functional endpoints, are robust and functional. Interpretation of this signal, however, is limited by the absence of per-cell normalization and orthogonal ATP/flux assays. Based on the set of assessed functional endpoints, we identified the 3dGRO freezing medium as the most promising candidate for organoid cryopreservation. Based on electrophysiological and Ca^2+^ imaging, apoptosis and immunofluorescent data, the organoids frozen in this medium were closer to controls than those frozen in the other two media tested. They did not have significant changes in neurite outgrowth and had the same levels of cleaved Caspase 3 as incubator controls. It is a commercial product that contains commonly used, relatively inexpensive, and readily accessible reagents, allowing other researchers worldwide to apply the protocol. This method facilitates the transport of ready-made brain organoids, increasing their accessibility to numerous research and pharma laboratories worldwide and enabling cross-laboratory research and harmonization.

In summary, we developed and tested a protocol for brain organoid cryopreservation. Based on the assessed freezing time points and endpoints, we concluded that freezing at an earlier stage of differentiation is better for overall recovery of functionality. Also, a recovery period is needed before organoids can be used, which can be assessed by organoids’ shape, size, and functionality. In our study, four weeks of recovery were enough to have functional organoids. Freezing enables the creation of organoids on demand from diverse donors, shortens culture time, and avoids laborious iPSC handling and differentiation. This provides a possibility for long-term storage and accelerates worldwide collaborations and global harmonization.

### Limitations

Although brain organoids successfully preserve several key aspects of human brain development and function, they still lack cellular complexity, including functional vascularization. Additionally, the present study focused on a single iPSC line; therefore, validation using multiple donor-derived lines across different protocols will be necessary to confirm the reproducibility of the findings. Furthermore, we did not quantify the fraction of organoids that survive, and recover function after freeze–thaw. Survival yield and recovery fractions will be needed to define banking throughput and release criteria.

## MATERIALS AND METHODS

### iPSCs culture

The commercially available iPSC line (NIBSC-8, National Institute for Biological Standards and Control (NIBSC), UK) was cultured feeder-free on Vitronectin XF (Gibco, ThermoFisher Scientific) in a chemically defined medium (mTESR-Plus, STEMCELL Technologies) at 5% O2, 5% CO2, and 37°C. Every 3 to 5 days, cells were passaged 1:10 with an enzyme-free passaging reagent (GDCR, Gentle Cell Dissociation Reagent, STEMCELL Technologies), and medium was changed every other day. Cells were mycoplasma-free with a normal female karyotype and were authenticated using STR profiling^31^.

### NPCs culture and expansion

iPSCs were seeded in mTESR-Plus medium as single cells, supplemented with Rock Inhibitor Y27632 (RI), and cultured at 5% O2, 5% CO2, and 37°C. After 24 hours, the medium was replaced with neural induction medium (49 mL of Neurobasal Medium (ThermoFisher Scientific) and 1 mL of Neural Induction Supplement (ThermoFisher Scientific)), and the cells were cultivated at 5% O2, 5% CO2, and 37°C for an additional six days. The medium was replaced every other day, and cellular morphology and induction were evaluated microscopically at 10x magnification. After seven days of neural induction, NPCs were expanded for four passages in the neural expansion medium: 24.5 mL of Neurobasal Medium, 24.5 mL of Advanced DMEM/F12 (ThermoFisher Scientific), and 1 mL of Neural Induction Supplement. For cryopreservation, NPC Freezing medium (StemCell Technologies) was used. To enhance survival, five μM RI was added to the medium at each passage for the first four passages. After the fifth passage, NPC were cultured in a normoxic incubator with 20% oxygen without RI. Medium was changed every 48 hours. NPCs were banked at passage 5-7. NPCs at passage 13 were used for the subsequent brain organoid differentiations and experiments performed in this study.

### 3D organoid formation

The differentiation protocol was adapted from Romero et al^31^. Briefly, after the NPCs reached 80-100% confluency, they were washed once with PBS and treated with Gentle Cell Dissociation Reagent (GCDR) for 4-5 min. GCDR was removed and cells resuspended in fresh neural expansion medium, scraped, and dissociated to a single cell suspension. 2.0×106 cells were plated in 3 mL of complete neural expansion medium per well in a clear 6-well plate. Cells were cultured at 37°C in 20% oxygen, 5% CO2 under constant gyratory shaking (88 rpm, 19 mm orbit). After 48 hours of culture, the medium is changed from the complete neural expansion medium to the neural differentiation medium: Neurobasal Plus, 1x B-27™ Plus Supplement (ThermoFisher Scientific), 1x Penicillin-Streptomycin-Glutamine (ThermoFisher), 1x GlutaMAX (ThermoFisher Scientific), 10 ng/mL GDNF (GeminiBio), and 10ng/mL BDNF (GeminiBio). Medium was changed every 2 to 3 days. Immune-competent organoids were generated following the modified Rittenhouse et al protocol^26^.

### Cryopreservation of brain organoids

For cryopreservation, we used three different freezing media: mFreSR (STEMCELL Technologies), CrystorCS10 (HemaCare), and 3dGRO (Millipore Sigma). For all the experiments, organoids were frozen at two time points of differentiation: 2 and 6 weeks. Before freezing, an isopropanol-based, controlled-rate freezing container (Mr. Frosty™, Thermo Fisher Scientific) was pre-cooled at 4°C for at least 30 min. Brain organoids were collected from 6-well plates and transferred to the pre-cooled cryopreservation vial using a 2 mL pipette – one well per vial, equivalent of 200-300 organoids. After organoids were precipitated, the medium was removed, and 1 mL of freezing medium was gently added to ensure the brain organoids were not clumped. Cryo-vials were transferred to Mr. Frosty and cooled at approximately 1 °C per minute to –80°C overnight, after which the cells were transferred into liquid nitrogen. Organoids that remained in the incubator and underwent routine culture served as a control for all assays. For the purpose of the experiments described here, organoids were frozen for 48 hours.

### Organoid defrosting protocol

Brain organoids were removed from liquid nitrogen and thawed rapidly in a 37 °C water bath, keeping an eye on the tube until only a small piece of ice remained. The organoids were then gently mixed with 9 mL Knockout Medium. After organoids sedimented, Knockout Medium was removed and organoids were gently resuspended in 3 mL of differentiation medium supplemented with 10 μM ROCK Inhibitor Y27632, incubated overnight on a shaker at 37°C in 20% oxygen, 5% CO2. Next day, the medium was exchanged to remove RI, and organoids were cultured routinely with medium change every 2-3 days. All endpoint assays were performed either 24 hours after defrosting or at 2 and 4 weeks after defrosting, indicating recovery periods of 24 hours and 2 and 4 weeks, respectively.

### Organoid size quantification

Following thawing, organoids were maintained in neural differentiation medium and imaged after 24 hour, 2 weeks, and 4 weeks of recovery. Brightfield images were acquired under identical imaging conditions, and organoids diameter was quantified using ImageJ. The maximum diameter of each organoid was measured and compared between frozen and control groups at each recovery time point to evaluate the impact of freezing medium. Statistical significance was determined by comparing each frozen condition with the corresponding control group, and data are presented as individual values with Mean±SD.

### Resazurin Reduction Assay

Ten organoids per condition were transferred to individual wells of a 96-well plate containing 100 μL of differentiation medium. Resazurin (10 μL; 1 mg/mL stock) was added to achieve a final concentration of 100 μg/mL, and the plate was incubated for 3 hours at 37°C and 5% CO_2_. Resazurin solution without organoids served as a blank to account for background fluorescence. Fluorescence was measured at 590 nm, and percent cell viability was calculated by comparing relative fluorescence units (RFU) from frozen samples to incubator controls using the following formula:

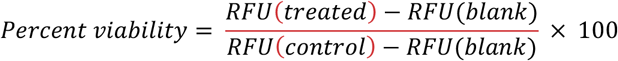

The measurements were done in triplicates. For each condition, three biological replicates were used, and within each biological replicate, three technical replicates were plated (a total of 9 wells per condition, with 10 organoids per well).

### Neurite outgrowth quantification

24h prior to the collection of samples, five organoids were plated per well in a 24-well glass-bottom black plate coated with PLO/laminin. After 2–3 days of incubation, when clear neurite outgrowth was observed, the organoids were fixed in 2% paraformaldehyde (PFA) for 1 h at 4°C, washed, permeabilized with 0.15% saponin, and blocked with 10% goat serum plus 1% Bovine Serum Albumin (BSA). Then the neurites were visualized by immunostaining with a β-III-tubulin antibody, while astrocytes were visualized with a GFAP antibody and their corresponding secondary antibody (Supplemental Table 1), and imaged with the Keyence BZ-X800. The density of neurites and glia processes was quantified as the number of intersections per distance from the organoid edge and presented as the area under the curve (AUC). The distribution’s normality was verified using the Shapiro-Wilk test, and statistical significance was determined using the one-way ANOVA test and Bonferroni’s multiple comparisons test (p < 0.05).

### MitoTracker assay

To assess mitochondrial membrane potential, brain organoids were stained with MitoTracker Red CMXRos. Briefly, 20 organoids per condition were transferred into a 24-well plate containing 500 μL neural differentiation medium supplemented with 1 μM MitoTracker. One mM MitoTracker stock solution was prepared in DMSO. Samples were incubated for 45 min at 37°C with 5% CO_2_ protected from light. After staining, organoids were washed for 1 hour with fresh differentiation medium followed by two PBS washes. Sample were fix with 4% PFA for 60 min at 4°C, mounted on glass slides, and imaged using Zeiss LSM710 confocal microscope under consistent settings across the samples. Mitochondrial membrane potential was quantified in ImageJ by measuring the mean gray value of MitoTracker fluorescence from 3 organoids per condition.

### Immunostaining

For immunostaining, at least 10 brain organoids were collected, fixed with 4% PFA for 1 h at 4°C, permeabilized in 0.1% Triton X-100 diluted in PBS for 30 min at 4 °C., and blocked with 100% BlockAid for 30 min at 4°C. After the blocking, organoids were stained with primary antibodies diluted in 10% BlockAid/0.1% Triton/PBS solution overnight at 4°C on a shaker. Then, organoids were washed three times for 1 hour each with Organoid Washing Buffer (0.1% Triton X-100 and 0.5% BSA in 1xPBS, OWB) and incubated with secondary antibodies diluted in 10% BlockAid/0.1% Triton/PBS solution overnight at 4°C on a shaker. Then, organoids were washed twice with OWB for one hour each. For the third wash, the OWB was replaced with PBS/1 % BSA containing Hoechst 33342 (1:5000). Organoids were then mounted on a glass slide with Immomount and imaged with a Zeiss LSM710 confocal microscope at 20x and 63x magnification. At least three organoids per condition were imaged. Antibodies used are listed in the Supplemental Table 1.

### RT-qPCR

RNA was extracted using the Quick-RNATM Microprep Kit (Zymo Research). RNA purity and concentration were quantified with the NanoDropTM 2000c Spectrophotometer (ThermoFisher Scientific). 500 ng of RNA were collected and reverse transcribed using the M-MLV Reverse Transcriptase (Promega). Gene expression was analyzed by the SYBR Green assay (ThermoFisher Scientific) on the 7500 Fast Real-Time PCR system (Applied Biosystems). Gene expression was normalized to the housekeeping gene, ACTINB. Fold change was calculated using 2^-ΔΔCt^. The primer sequence is listed in Supplemental Table 2.

### Calcium imaging

Calcium imaging and data analysis were conducted following our recently published protocol (Alam El Din et al., 2025). Brain organoids were incubated with 10 μM Fluo-4 AM (Tocris) and 0.5% Pluronic Acid (Invitrogen) for 2 hours at 37 °C and 5% CO_2_ in the dark, without shaking. Then wash three times with neural differentiation medium. After washing, ganoids were transferred to a glass-bottom 24-well plate and imaged using an Olympus FV3000-RS confocal microscope equipped with a resonant scanner with a speed of 12.5 frames per second for 6 minutes. The temperature and CO_2_ levels were maintained at 37 °C and 5% respectively, during the imaging. Fluorescence intensity over time was quantified using the ROI Manager in ImageJ and analyzed with a custom Python script to generate ΔF/F plots (https://github.com/organoid-intelligence/bMPS_analysis_tools).

### High-density microelectrode array recordings

The extracellular field potential of the brain organoids was recorded using the MaxTwo Multi-well MEA platform as described in Alam El Din et al., 2025. Briefly, organoids were plated on pre-coated MEA plates on day 42 of differentiation. Spontaneous electrical activity was recorded every 3 days for up to 31 days. MaxWell Biosystems 6-well HD-MEA chips were coated for 1 hour at 37 °C, 5% CO_2_ with 0.07% poly(ethyleneimine) in 1× borate buffer (Sigma-Aldrich, ThermoFisher), washed ×3 with water, and dried for 1 hour. Chips were then coated with 0.04 mg/mL mouse laminin (Sigma-Aldrich) in brain differentiation medium for 1 hour under the same conditions. After removing laminin, 2-week organoids were placed on the MEA in differentiation medium. Recordings were taken from day 3 to 31 (DOM 0 = plating day) for organoids frozen at week 2 and from day 7 to day 43 for the organoids frozen at week 6. MaxLab Live v22.2.6 (MaxWell Biosystems, Switzerland) was used to collect data, and MaxLab Live v25.1.8.1 was used to analyze percent active area and network spiking and bursting metrics. Recordings used a 512× gain, 300 Hz high-pass filter, and 5.5 RMS mV threshold. A threshold of at least 100 spikes was determined to classify a “burst”. At least 3 wells per group per time point across two independent experiments were recorded and analyzed for organoids frozen at week 2, and one independent experiment for organoids frozen at week 6. Several organoids (up to 5) were plated per well, the number of organoids per well was not controlled as this experiment’s goal was to determine if freezing organoids still have spontaneous activity and not to compare activity levels between them. Therefore, we do not perform statistical comparisons among activity dynamics.

### Quantification and statistical analysis

In this study, we defined individual organoids as technical replicates, and one frozen vial of organoids as a biological replicate. An independent experiment was defined as an independent differentiation, where at least three vials of organoids were frozen per experiment. Data is represented as Median or Mean±SD or SEM, as denoted in the figure legends and method section for each data set. Statistical significance of size differences was assessed with One-Way ANOVA and Dunnett post-hoc test. Statistical differences in viability, neurite outgrowth, and glia migration, mitotracker assay, and caspase 3 assay were calculated with the Kruskal-Wallis test, with Dunn’s post-hoc test. Statistical significance for qRT-PCR was calculated with one-way ANOVA. The significance of the calcium imaging was assessed using a one-way ANOVA with Dunnett’s multiple comparisons test. ns – not significant, ** p< 0.01, *** p< 0.001, **** p< 0.00001.

## RESOURCE AVAILABILITY

### Lead contact

Requests for further information and resources should be directed to and will be fulfilled by the lead contact, Lena Smirnova.

### Materials availability

This study did not generate new unique reagents.

### Data and code availability

The calcium imaging code has been published previously in (Alam El Din et al., 2025) and deposited at https://github.com/organoid-intelligence/bMPS_analysis_tools and is publicly available.

## ACKNOWLEDGMENTS

We thank all members of the Center for Alternatives to Animal Testing for technical help and support. We thank the Integrated Imaging Center (NIH SIG award #1S10 OD020152-01A1), the Department of Neuroscience Multiphoton Imaging Core at Johns Hopkins University and the JHU SOM Microscope Facility. We thank George McNamara, PhD, the Ross Fluorescence Imaging Center (Johns Hopkins University), and the NIH shared instrumentation grant 1S10OD025244-01 for the use of the FV3000RS. D.M.A.E.D was supported by the National Institutes of Health (T32 ES007141) and the International Foundation for Ethical Research Graduate Fellowship.

## AUTHOR CONTRIBUTIONS

L.D., T.H., and L.S. designed and conceptualized the work, assembled figures, and wrote the manuscript. J.Z. cell culture, imaging, qPCR and data analysis, writing, methodology. D.A. Conceptualization, Methodology, Software, Validation, Formal analysis, Investigation, Writing. I.E.M.P. MEA recording and helping with staining. LS funding acquisition.

