## Supplemental Materials for "Cryopreservation of brain organoids - a tool for on-demand organoid banking"

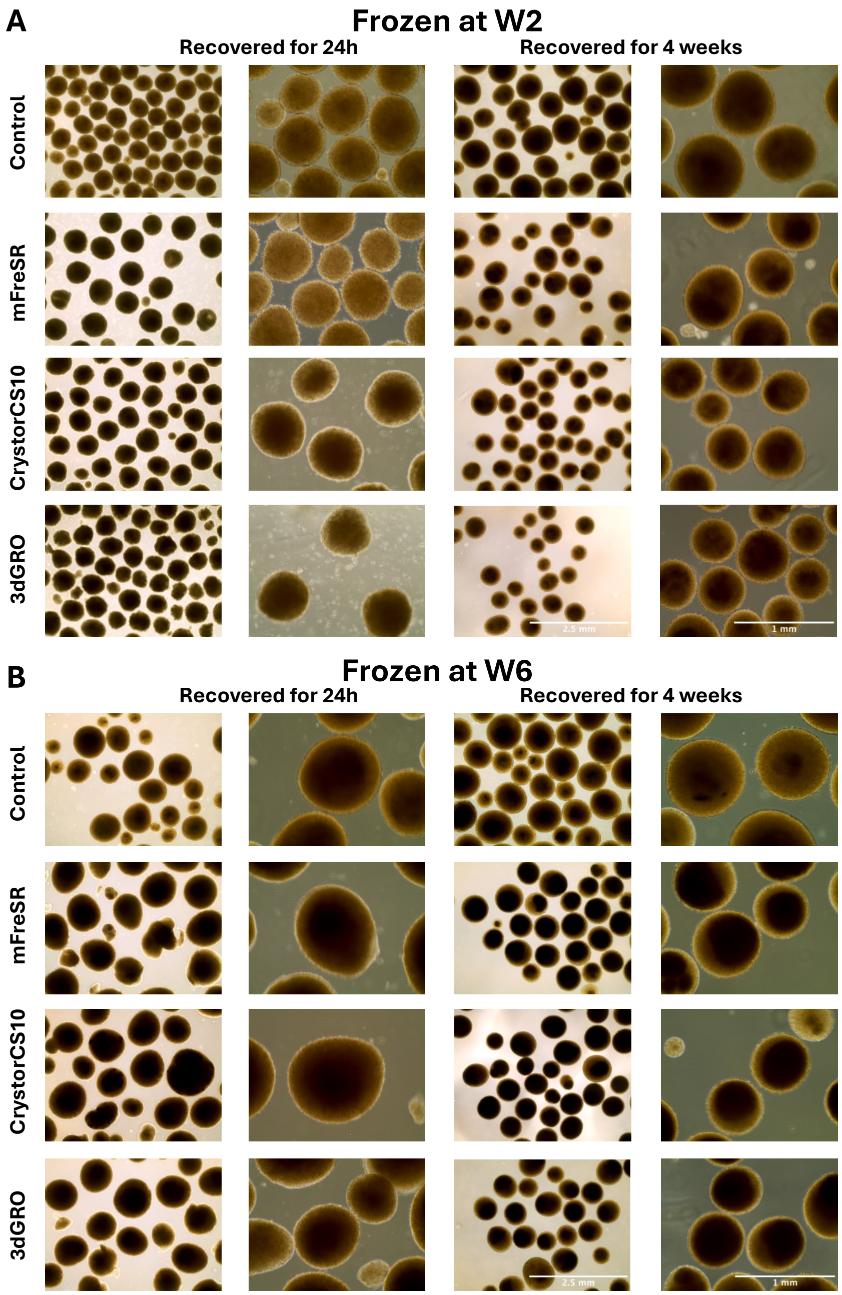

**Supplemental Figure S1. Morphology of cryopreserved organoids.**

Representative images of brain organoids, cryopreserved at (A) week 2 (W2) and (B) week 6 (W6) of differentiation using three different freezing media: mFreSR, CryostorCS10, 3dGRO. Age-matched organoids that remained in the incubator throughout the experiment served as a control. Bright field images show organoid shape and morphology 24 hours or 4 weeks after recovery. Scale bars are 2.5 and 1 mm.

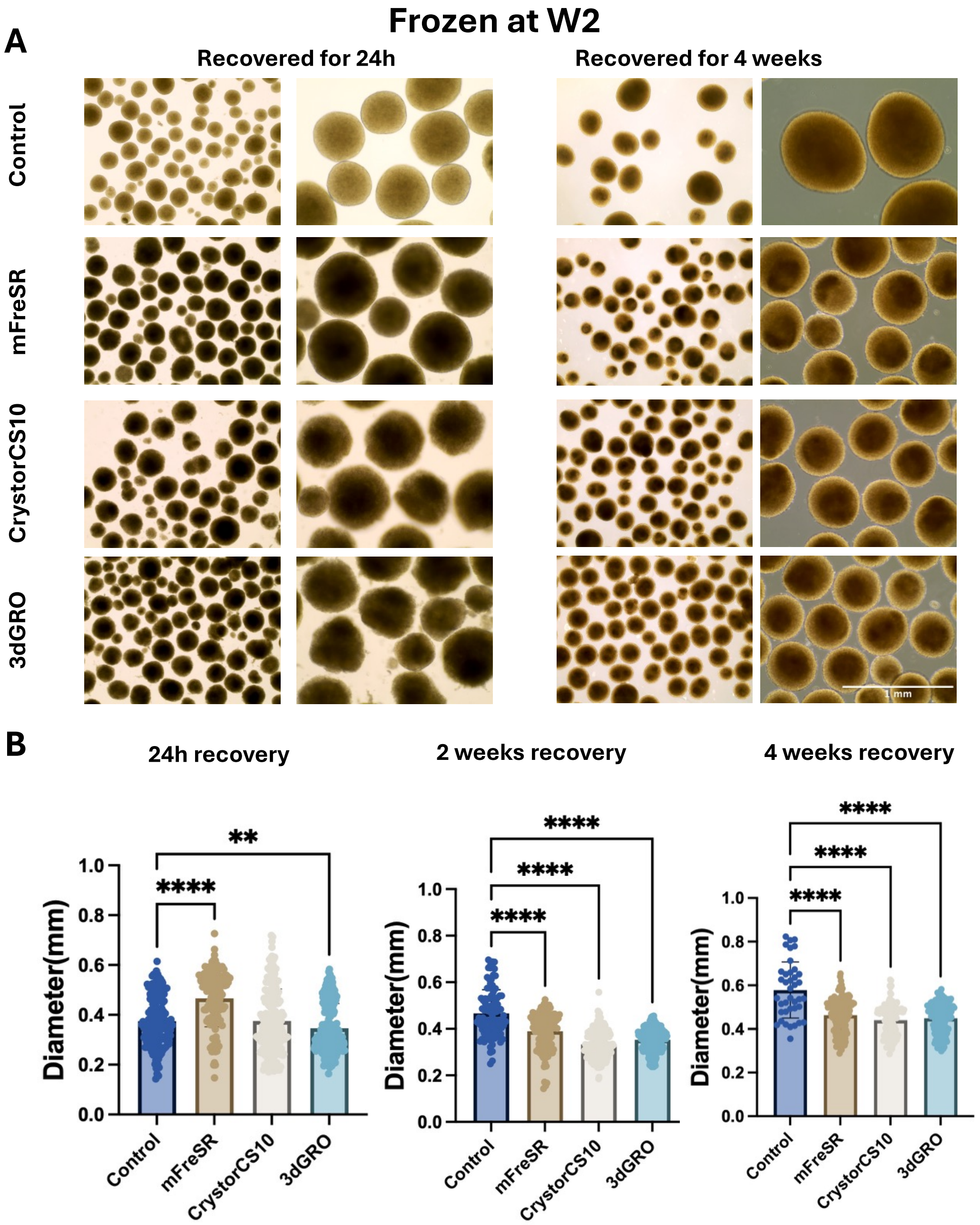

**Supplemental Figure S2. The morphology and quantification of brain organoid size frozen at week 2 in an additional independent experiment**

Organoids were frozen at 2 weeks (W2) and recovered for 24 hours, 2 or 4 weeks. Age-matched organoids that remained in the incubator throughout the experiment served as a control. (A) Representative images. (B) Diameter quantification. Data are presented as Mean ± SD from 3 biological replicates with at least 40 organoids measured in total for each condition and time point. One-way ANOVA with Dunnett's post-hoc test was used to assess statistical significance. ** $p<0.01$, **** $p<0.0001$.

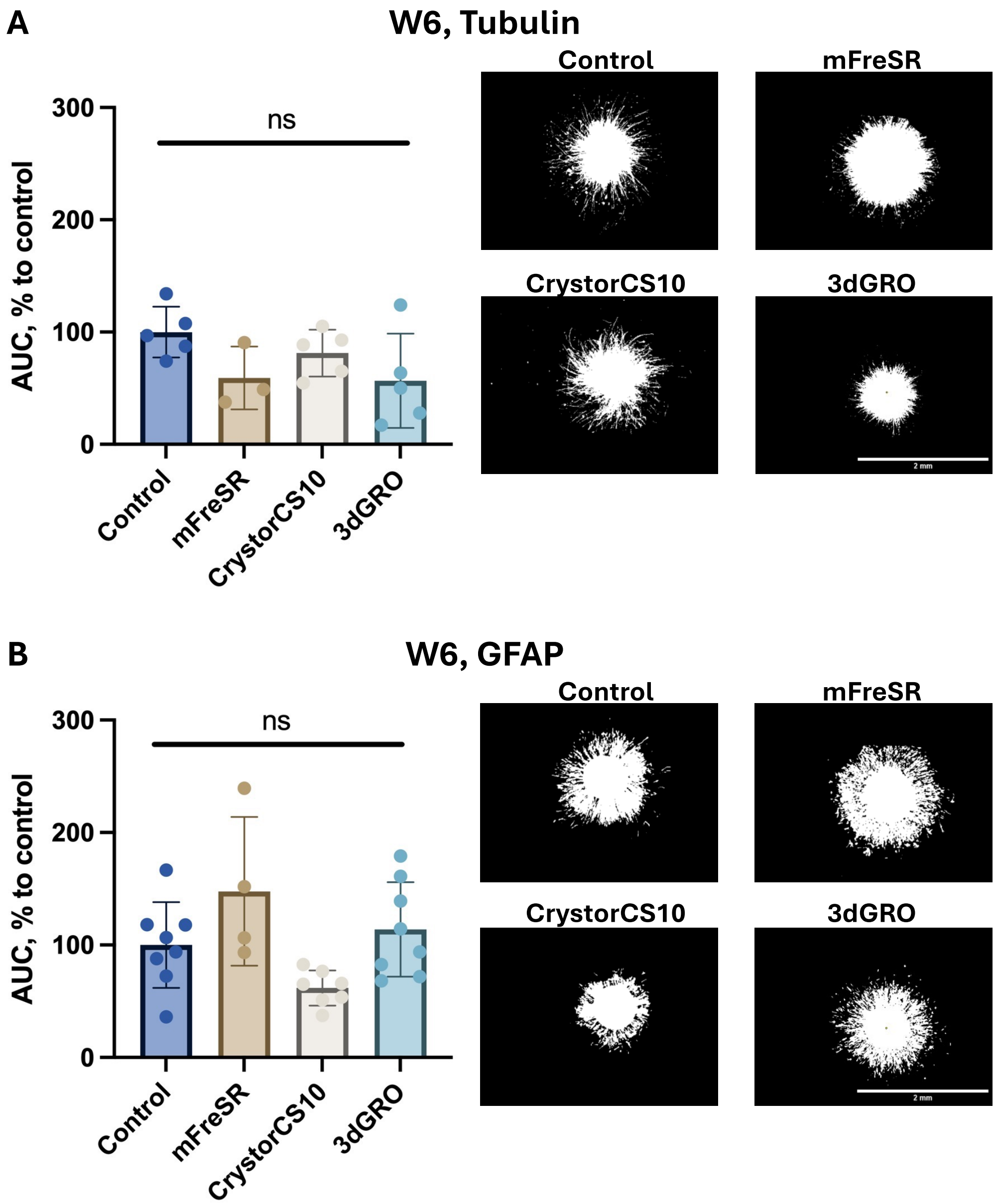

**Supplemental Figure S3. Neurite outgrowth and astrocyte migration in brain organoids frozen at week 6 and recovered for 4 weeks.**

Area Under the Curve (AUC) from Sholl analysis of (A) neurite outgrowth (β-III-Tubulin) and (B) astrocyte migration (GFAP), and representative organoids after transformation to binary images are shown. The area under the curve (AUC) was calculated for each condition, and each graph represents the Mean ± SD from 3-7 organoids per independent experiment, with one experiment for Tubulin and two experiments for GFAP. Kruskal-Wallis test with Dunn's post-hoc test was used to assess statistical significance. ns - not significant, *p<0.05, ***p<0.001.

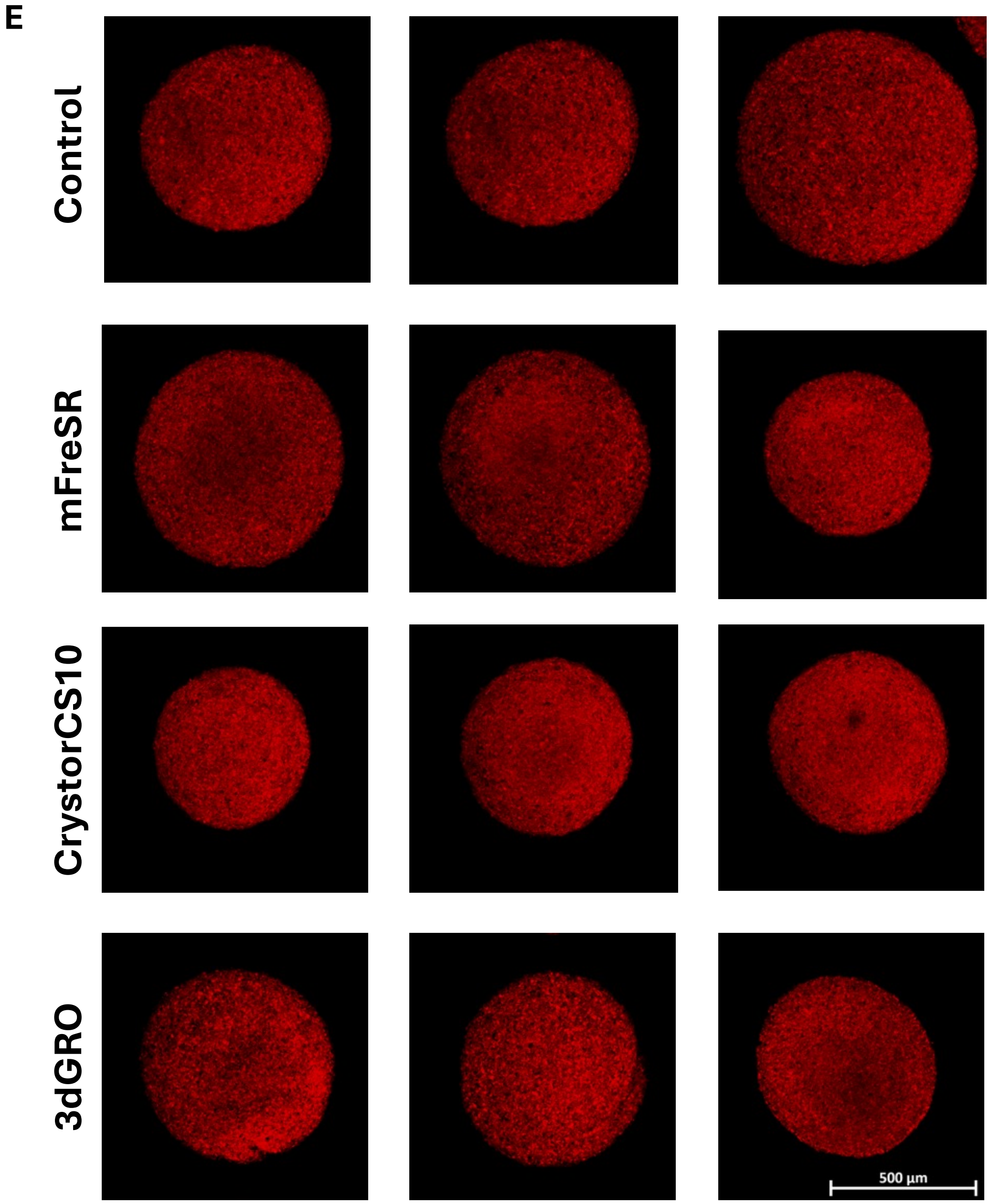

**Supplemental Figure S4. Fluorescent images of the Mitochondria membrane potential assay, which were used for quantification in Figure 6C.**

Representative fluorescent signal intensity reflects mitochondrial membrane potential, with brighter signal indicating higher mitochondrial activity.

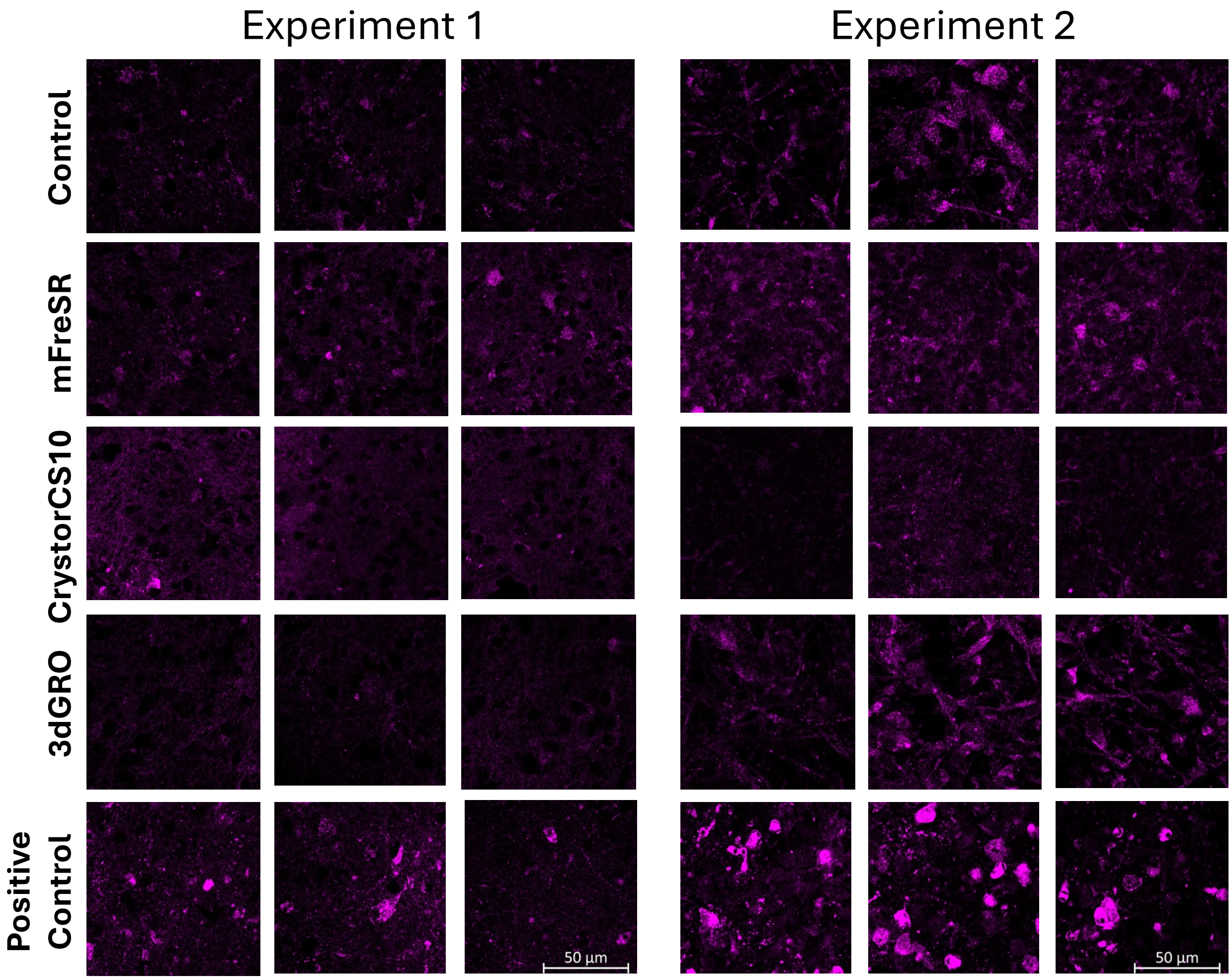

**Supplemental Figure S5. Confocal immunofluorescence images of cleaved caspase 3 used for quantification analysis, shown in Figure 6D.**

A positive control confirms assay performance. Six organoids per condition were assessed in two independent experiments.

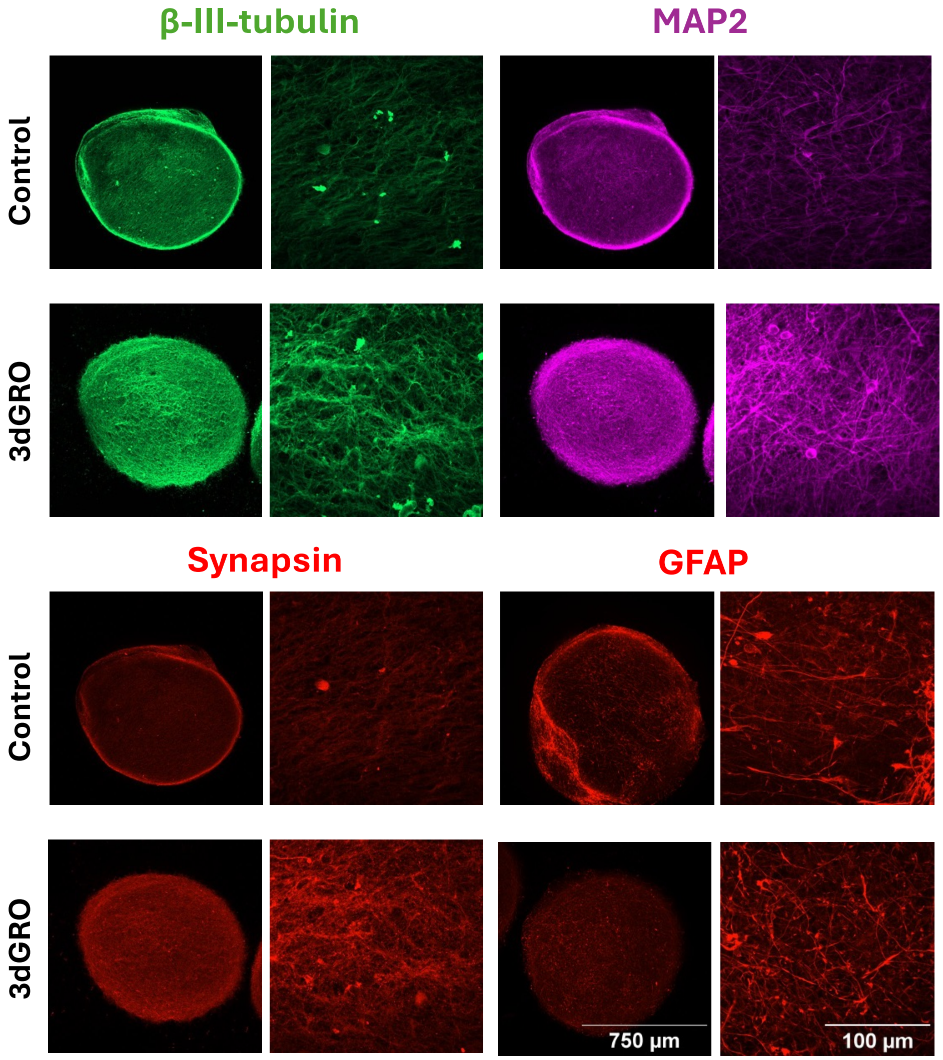

**Supplemental Figure S6.** Expression of neural markers (β-III-tubulin, GFAP, synapsin, and MAP2) in immune-competent microglia-containing organoids cryopreserved at week 2 and then recovered for 4 weeks.

**Supplemental Table 1. List of antibodies used for immunofluorescence**

| Antigen | Host Species | Vendor | Catalog No. | Dilution used |
|---|---|---|---|---|
| **B-TUBULIN** | Mouse | Milpore sigma | AB9354 | 1:1500 |
| **MAP2** | Chicken | Invitrogen | PA1-10005 | 1:5000 |
| **Synapsin** | Rabbit | Millipore | S193-10UG | 1:500 |
| **CD68** | Mouse | Invitrogen | 14-0688-37 | 1:100 |
| **PU.1** | Mouse | Novus biologicals | NBP2-75738-20ug | 1:500 |
| **IBA1** | Chicken | aveslabs | IBA1-0100 | 1:400 |
| **GFAP** | Rabbit | Agilent | GA524 | 1:250 |

**Supplemental Table 2. List of primers used for RT-qPCR**

| Gene | Name | Sequence (5'>3") |
|---|---|---|
| GFAP | GFAP_Forward | F: AATGGCGTCCAGCAACATGC |
| | GFAP_Reverse | R: CAGCAGCGTCTGTCAGGTCT |
| MAP2 | MAP2_Forward | F: GCGGAAAACCACAGCAGCA |
| | MAP2_Reverse | R: CCGAGGAGGGAGAATGGAGG |
| RBFOX3 | RBFOX3_Forward | F: TGAGAACTCCCGGCTGCAAA |
| | RBFOX3_Reverse | R: CCCTCTGGGGTCCTAGGGAA |
| NEST | NEST_Forward | F: TGGCCACGTACAGGACCCTC |
| | NEST_Reverse | R: GAGCAAAGATCCAAGACGCCG |
| SYN1 | SYN1_Forward | F: TGGCCACGTACAGGACCCTC |
| | SYN1_Reverse | R: GAGCAAAGATCCAAGACGCCG |
| ACTB | ACTB_Forward | F: 5'-CCTCGCCTTTGCCGATCC-3' |
| | ACTB_Reverse | R: 5'-CGCGGCGATATCATCATCCAT-3' |

**KEY RESOURCES**

| REAGENT OR RESOURCE | SOURCE | IDENTIFIER |
|---|---|---|
| Antibodies | | |
| Anti-β-Tubulin III Antibody | Millipore Sigma | Cat #AB9354 |
| MAP2 | Invitrogen | Cat #PA1-10005 |
| Anti-Synapsin I | Millipore Sigma | Cat #S193-10UG |
| Anti-Glial Fibrillary Acidic Protein | Agilent | Cat #GA524 |
| Anti-Neurofilament 200 antibody | Millipore Sigma | Cat #N4142 |
| Anti-IBA1/AIF1 Antibody | aveslabs | Cat #IBA1-0100 |
| PU.1/Spi-1 Antibody | Novus biologicals | Cat # NBP2-75738-20ug |
| CD68 Monoclonal Antibody (KP1) | Invitrogen | Cat # 14-0688-37 |
| Cleaved caspase-3 (Asp175) Antibody | Cell signaling Technology | Cat #9661S |
| Alexa Fluor 488 Goat Anti-Mouse | Invitrogen | Cat #A11001 |
| Alexa Fluor 568 Goat Anti-Rabbit | Invitrogen | Cat #A11011 |
| Alexa Fluor 647 Goat Anti-Chicken | Invitrogen | Cat #A21244 |
| Alexa Fluor 647 Goat Anti-Rabbit | Invitrogen | Cat #A21244 |
| Biological samples | | |
| Human Induced Pluripotent Stem Cells (iPSC) | See Experimental Models: Cell lines | See Experimental Models: Cell lines |
| Human Neural Progenitor Cells (NPC) | This paper | N/A |
| Human Brain Organoids derived from NPC | This paper | N/A |
| Chemicals, peptides, and recombinant proteins | | |
| Fluo-4 AM | TOCRIS | Cat #6255 |
| Hoechst | Invitrogen | Cat #H3570 |

| | | |
|---|---|---|
| mTeSR™ Plus | STEMCELL Technologies | Cat #100-0276 |
| Vitronectin (VTN-N) Recombinant Human Protein, Truncated | Gibco | Cat #A14700 |
| B-27™ Plus Neuronal Culture System | Gibco | Cat #A3653401 |
| Human Glial Derived Neurotrophic Factor | GeminiBio | Cat #300-121P-100 |
| Human Brain Derived Neurotrophic Factor | GeminiBio | Cat #300-104P-100 |
| Y - 27632 | STEMCELL | Cat #72305 |
| Neurobasal™ Medium | Gibco | Cat #21103049 |
| KnockOut™ DMEM/F-12 | Gibco | Cat #12660012 |
| PSC Neural Induction Medium | Gibco | Cat #A1647801 |
| GlutaMAX™ Supplement | Gibco | Cat #35050061 |
| Quality Biological Inc PBS (10X), pH 7.4, 1000 mL | Quality Biological Inc | Cat #119069131 |
| Bovine Serum Albumin | Sigma Aldrich | Cat #A9418 |
| Gentle Cell Dissociation Reagent | STEMCELL Technologies | Cat #100-0485 |
| Immu-Mount | Epredia | Cat #9990402 |
| Penicillin-Streptomycin-Glutamine (100X) | Gibco | Cat #10378016 |
| mFreSR™ | STEMCELL Technologies | Cat #5855 |
| CryoStor® CS10 | STEMCELL Technologies | Cat #07959 |
| 3dGRO® Organoid Freeze Medium | Sigma Aldrich | Cat #SCM301 |

| | | |
|---|---|---|
| STEMdiff™ Neural Progenitor Freezing Medium | STEMCELL Technologies | Cat #05838 |
| Paraformaldehyde | Sigma Aldrich | Cat #P6148-500G |
| BlockAid™ Blocking Solution | Invitrogen | Cat #B10710 |
| Saponin | EMD Millipore | Cat #2960459 |
| Triton X-100 | SIGMA | Cat #X100-500ML |
| Corning™ Matrigel™ hESC-Qualified Matrix | Corning | Cat #354277 |
| Resazurin sodium salt | SIGMA | Cat #R7017-1G |
| Trypan Blue stain 0.4% | Invitrogen | Cat #2685419 |
| M-MLV Reverse Transcriptase | Promega | Cat #M1701 |
| RNaseOUT™ Recombinant Ribonuclease Inhibitor | Invitrogen | Cat #10777019 |
| DNase I Set (RNase-free) | ZYMO RESEARCH | Cat #E1012 |
| M-MLV RT 5X Buffer | Promega | Cat #M531A |
| Random Primers | Promega | Cat #1181 |
| STEMdiff™ Hematopoietic Kit | STEMCELL Technologies | Cat #05310 |
| Pluronic™ F-127 (20% Solution in DMSO) | Invitrogen | Cat #P3000MP |
| Critical commercial assays | | |
| Quick-RNA Microprep Kit | ZYMO RESEARCH | Cat #1051 |
| Experimental models: Cell lines | | |
| Human: NIBSC8 (N8) Experimental models: Cell lines | National Institute for Biological Standards and Control, NIBSC (NIBSC), UK | N/A |
| Oligonucleotides | | |

| | | |
|---|---|---|
| Primers for RT-qPCR Experiments | This paper | See Table S2 for Primers used in RT-qPCR Experiments |
| Software and algorithms | | |
| FIJI(Version 1.54p) | | https://imagej.net/software/fiji/ |
| GraphPad Prism 9.0 | GraphPad Software Inc. | https://www.graphpad.com/ |
| Python | Alam El Din et a. 2025 | https://github.com/organoid-intelligence/bMPS_analysis_tools |
| Other | | |
| Zeiss LSM 700 Microscope | Zeiss | |
| Falcon® 6-well Clear Flat Bottom Not Treated Cell Multiwell Culture Plate | Falcon | 351146 |
| 24 Well glass bottom plate with high performance #1.5 cover glass | Cellvis | Cat #P24-1.5H-N |
| 96 Well glass bottom plate with high performance #1.5 cover glass | Cellvis | Cat #P96-1.5H-N |
| CryoFreeze® Cryogenic Vials: Internally-threaded cryo tubes, with O-ring cap seal | Avantor | Cat #77093-430 |
| 10µl Reach Olympus Ergonomic Pipet Tips Low Binding | Olympus | 24-121RL |
| 200µl Reach Olympus Ergonomic Pipet Tips Low Binding | Olympus | 24-150RL |
| 1000µL Reach Olympus Ergonomic Pipet Tips Low Binding | Olympus | 24-165RL |

| | | |
|---|---|---|
| Keyence All-in-One Fluorescence Microscope | Keyence | BZ-X1000 |
| 5mL Serological Pipettes, Paper/Plastic Peel, Individually Wrapped, Pack of 100 | ThermoFisher | Cat #170355N |
| 10mL Serological Pipettes, Paper/Plastic Peel, Individually Wrapped, Pack of 100 | ThermoFisher | Cat #170356N |
| Nunc™ Serological Pipettes (25 mL) Individually Wrapped, paper/plastic peel packaging, plugged | ThermoFisher | Cat #170357N |
| Fisherbrand™ Sure One™ Filtered Pipette Tips | ThermoFisher | Cat #02-707-404 |
| Eppendorf® ep Dualfilter T.I.P.S | Eppendorf® | EP0030078519 |
| ART™ Barrier Pipette Tips in Lift-off Lid Rack | ThermoFisher | Cat #2069 |
| Fisherbrand™ SureOne™ Low Retention Filtered Pipette Tips | ThermoFisher | Cat #02707003 |
| Olympus FV3000-RS confocal | OLYMPUS | |
| MaxTwo 6-Well-Plate | Maxwell Biosystems | MX2-S-6W |
| MEA machine | Maxwell Biosystems | |
| PolysineTM Slides | epredia | Cat #P4981-001 |
| Microscope cover glass | Globe scientific | Cat #1415-15 |
| CellCarrier Spheroid ULA 96-well Microplates | revvity | Cat #6055330 |
| VWR CO2 Incubator symphony 5.3 A | VWR | Cat #50111308B |
